# Local translational program at the muscle-tendon junction endows domain identity in muscle syncytia

**DOI:** 10.64898/2026.01.01.697307

**Authors:** Juliana de Carvalho Neves, Nour El Khazen, Coalesco Smith, Vedran Franke, Sakulrat Mankhong, Edgar Jauliac, Laura Yedigaryan, Elise Lefebvre, Altuna Akalins, Pascal Maire, Minchul Kim

## Abstract

How cells establish specialized subdomains is a fundamental question in cell and tissue biology. Skeletal muscle fibers, among the largest cells in the body, are multinucleated and form distinct regions such as the neuromuscular and myotendinous junctions (MTJ), the latter forming a critical interface between muscle and tendon that transmits contractile force. While transcriptional heterogeneity among myonuclei has been described, whether local translation contributes to domain identity remains unknown, largely due to the lack of tools for domain-specific manipulation. Here, we introduce MTJ-AAV, a viral system that enables selective genetic targeting of MTJ myonuclei. This approach allowed MTJ-specific ribosome tagging and revealed extensive translational regulation underlying MTJ biology and its remodeling during exercise. Interestingly, untranslated regions of these transcripts were sufficient to control regionalized translation. Notably, the KLF-family transcription factors emerged as translationally upregulated targets at the MTJ, where they drive local gene expression. Our findings establish local translation as a key layer of subcellular specialization and provide a versatile toolkit for dissecting spatial molecular regulation within muscle syncytia.

## Introduction

To precisely coordinate diverse biochemical activities and biological functions, cells must establish specialized subcellular domains. How these domains form, how they are maintained and remodeled remain a fundamental question in cell biology. Skeletal muscle cells, or myofibers, provide a unique model to study this process because of their exceptionally large cytoplasm, which contains hundreds to thousands of nuclei. Different myofiber regions interact with distinct external cues, giving rise to specialized functional domains. For example, myofiber central regions interact with motor neurons, forming the neuromuscular junction (*1, 2*), while their ends attach to tendons, creating the myotendinous junction (MTJ) (*3, 4*). The MTJ plays a critical role in dissipating the mechanical forces generated during contraction and transmitting them to tendons. Consequently, the MTJ is highly susceptible to injury, and its dysfunction leads to muscle rupture (*5*). Understanding how these muscle domains are regulated not only advances our knowledge of myology and disease, but also provides broader insights into how cells spatially and functionally organize their intracellular space.

Previous studies using single-nucleus RNA sequencing (snRNA-Seq) have shown that muscle nuclei at specialized domains adopt distinct gene expression programs, conceptually resembling cellular differentiation in multicellular tissues (*6–9*). These studies highlight transcriptional control as an important mechanism for domain formation. However, the contribution of post-transcriptional regulation remains largely unexplored. Translational regulation, the control of protein synthesis from existing mRNA, has emerged as a central mechanism across diverse biological contexts, including gametogenesis, neurodevelopment, cancer, and inflammatory signaling (*10–14*). In muscle, translational profiling has thus far been limited to whole myofibers (*15–17*), leaving little insight into how localized protein synthesis supports muscle domain biology. Interestingly, although the MTJ undergoes structural remodeling during exercise and aging (*18, 19*), snRNA-Seq studies have not detected major transcriptomic changes in MTJ myonuclei (*7, 20*). This discrepancy suggests that post-transcriptional regulation, particularly translation, may play a key role in these adaptations.

Conventional genetic approaches in the muscle field lack spatial resolution and target all myonuclei indiscriminately. The distinct lack of genetic tools that can manipulate specific domains is the major roadblock preventing the field from studying regionalized molecular events in myofibers. To overcome this limitation, we reasoned that promoters of MTJ-specific genes identified from snRNA-Seq datasets could be leveraged to drive gene expression specifically within this domain.

Here, we develop a versatile adeno-associated virus (AAV) system that enables expression of genetic tools specifically at the MTJ. Using this platform, we profile the local translational landscapes of the MTJ and uncover extensive translational regulation that becomes markedly amplified during endurance exercise. We further demonstrate that untranslated regions (UTRs) of MTJ-enriched transcripts are sufficient to confer localized translation and that KLF transcription factors are translationally upregulated to drive local gene expression. Together, these findings reveal a previously unrecognized layer of spatial regulation underlying this specialized muscle domain, and provide a powerful toolkit for dissecting subcellular regulation in a syncytium.

## Results

### Development of MTJ-AAV

To selectively target the MTJ, we focused on *Tigd4*, a highly specific marker of MTJ nuclei (*6*). We synthesized a fragment of the *Tigd4* promoter, spanning from the transcription start site to -3.3 kb, and cloned it into an AAV plasmid to drive the expression of super-fold GFP (sfGFP) fused with four SV40-derived nuclear localization signals (NLS). The 3.3 kb length was chosen to accommodate AAV genome size constraints. The resulting pAAV-Tigd4-NLS-sfGFP construct was packaged into the MyoAAV4a serotype (*21*) and intramuscularly injected into 10-week-old wild-type (WT) mice (Fig. 1A). As a control, we performed parallel experiments using the conventional *CK8* promoter, which is active in all myonuclei.

**Figure 1.**
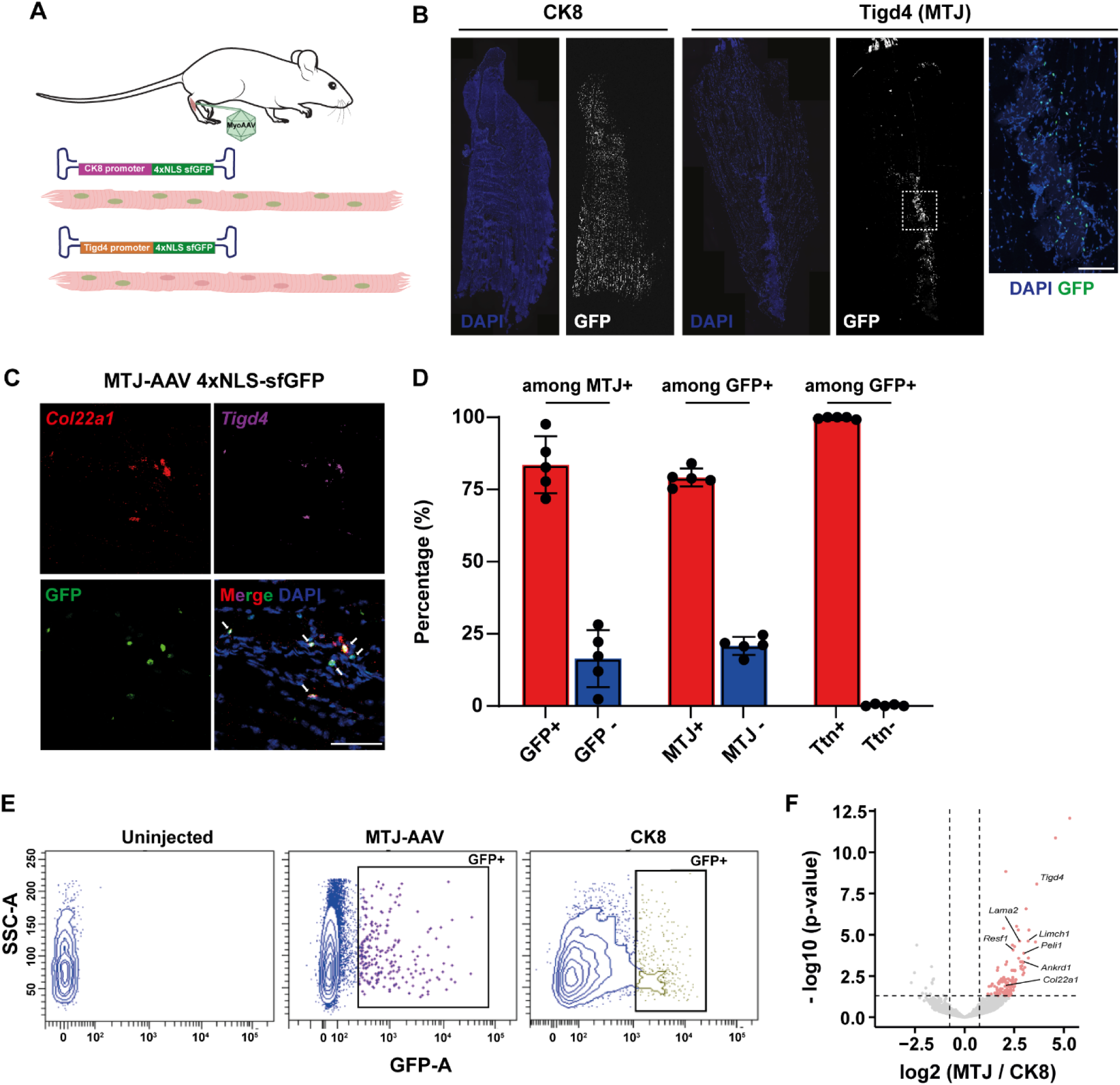
Development of MTJ-AAV. **(A)** Schematic of AAV constructs driven by *CK8* or *Tigd4* promoters and the expected expression pattern of the 4×NLS-sfGFP reporter. **(B)** Tile-scan images of tibialis anterior (TA) muscles injected with the AAVs shown in (A). The right panel shows a magnified view of the boxed region. **(C)** Combined GFP immunostaining and RNAscope detection of *Col22a1* and *Tigd4* transcripts. Arrows indicate nuclei co-expressing all three markers. **(D)** Quantification of labeling efficiency and specificity from experiments in (C) (n = 5). **(E)** Representative FACS plots of CK8- and Tigd4-NLS-sfGFP injected TA muscles. GFP+ nuclei were isolated and subjected to bulk RNA-Seq. **(F)** Heatmap showing genes enriched or depleted in GFP+ nuclei labeled by the *Tigd4* promoter relative to the *CK8* promoter. Selected MTJ marker genes are indicated. Scale bars, 100 µm.

As expected, *CK8* promoter-driven AAV broadly labeled the entire muscle tissue with NLS-sfGFP when *tibialis anterior* (TA) muscles were examined one-month post-injection. In contrast, the *Tigd4* promoter restricted reporter expression to the MTJ, identifiable by characteristic DAPI-stained structures penetrating into the muscle (Fig. 1B). Quantification of labeling efficiency using GFP immunostaining combined with RNAscope for the MTJ markers *Tigd4* and *Col22a1* revealed that approximately 80% of MTJ nuclei (defined by co-expression of *Tigd4* and *Col22a1*) were GFP-positive (Fig. 1C and 1D). Conversely, only ∼20% of GFP-positive nuclei were not MTJ nuclei, indicating high labeling specificity. Importantly, all GFP-positive nuclei were myonuclei as they also expressed *Ttn*, a pan-myonuclei marker (Fig. 1D), and located adjacent to the MTJ.

To confirm the molecular identity of the labeled nuclei via an orthogonal method, we isolated GFP-positive nuclei by FACS (Fig. 1E) and performed bulk RNA sequencing (Fig. 1F). The analysis showed that many representative MTJ markers, including *Tigd4, Col22a1*, *Resf1, Lama2* and *Ankrd1*, to be enriched in GFP-positive nuclei from the *Tigd4* promoter-driven AAV compared to those labeled by the *CK8* promoter (Fig. 1F). From here on, we refer to our new AAV system that uses *Tigd4* promoter as ‘MTJ-AAV’.

We next tested whether we could express other genetic tools at the MTJ as it will broaden the utility of MTJ-AAV. As such, we asked whether MTJ-restricted recombination could be achieved by delivering Cre recombinase. MTJ-AAV carrying Cre-NLS was intramuscularly injected into *Rosa26-LSL-H2B-GFP* reporter mice (Fig. 2A), resulting in GFP labeling around the MTJ (Fig. 2B). Quantification using MTJ markers showed a similar labeling efficiency of ∼75% (Fig. 2C). However, off-target labeling was higher in this model, with ∼40% of GFP-positive nuclei not corresponding to MTJ nuclei (Fig. 2C). Similar to our previous approach, all labelled nuclei were *Ttn*-positive, confirming their myonuclear identity.

**Figure 2.**
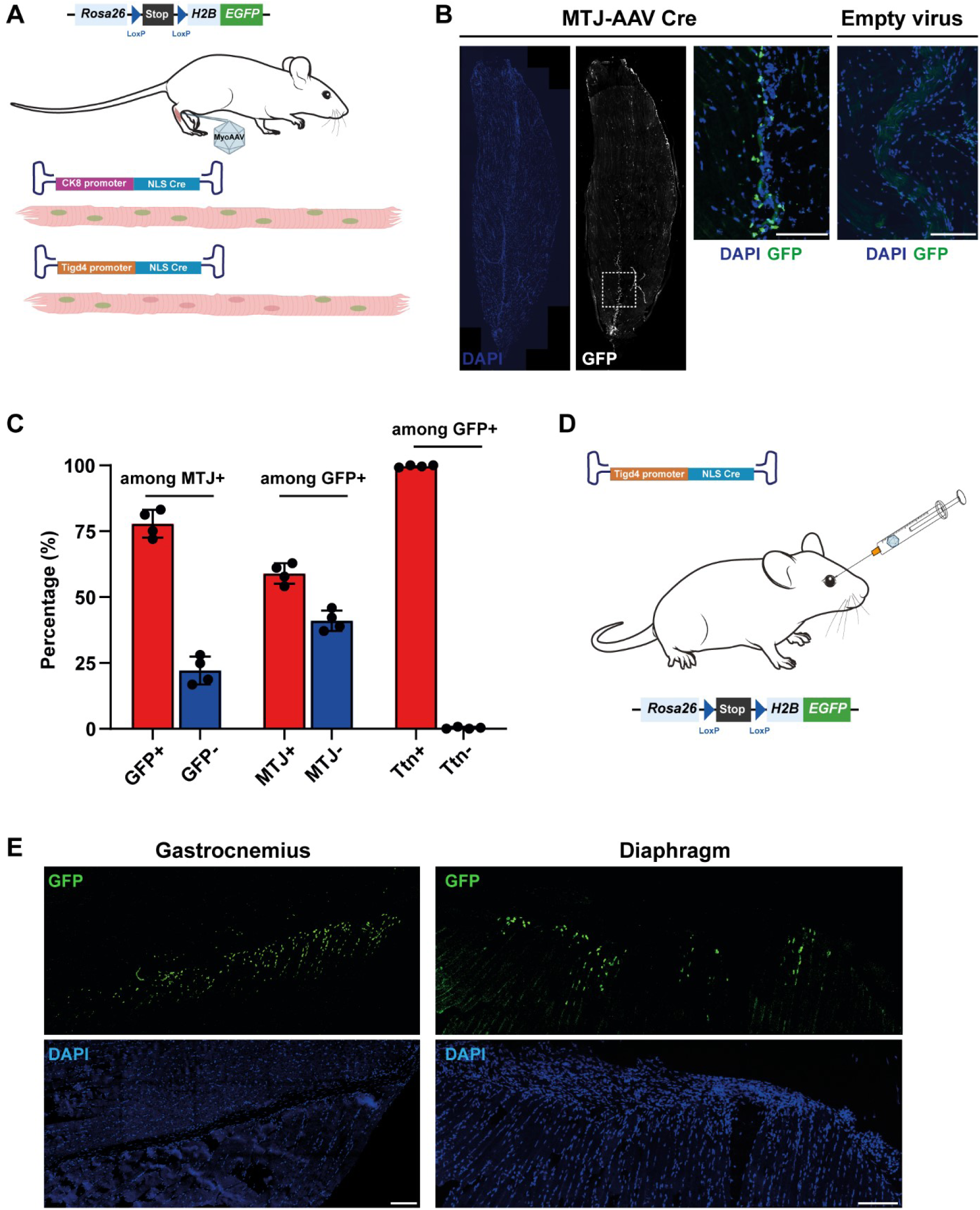
MTJ-AAV enables Cre-mediated recombination specifically at the MTJ. **(A)** Schematic of the AAV construct in which the *Tigd4* promoter drives expression of Cre-NLS. When injected into *Rosa26-LSL-H2B-GFP* reporter mice, Cre activity induces H2B-GFP labeling of MTJ nuclei. **(B)** Tile-scan images of TA muscles injected with MTJ-AAV-Cre-NLS. Reporter mice injected with an empty MyoAAV4a vector (containing only the *Tigd4* promoter without Cre) showed no H2B-GFP expression. **(C)** Quantification of labeling efficiency and specificity from experiments in (B) (n = 4). **(D)** Schematic of systemic delivery of MTJ-AAV-Cre-NLS into *Rosa26-LSL-H2B-GFP* reporter mice via intra-orbital injection. **(E)** Representative images showing MTJ-specific myonuclear labeling in the gastrocnemius and diaphragm muscles. Similar results were obtained in three independent animals. Scale bars, 100 µm.

To investigate why there was higher off-target labelling with Cre compared with NLS-sfGFP, we first tested whether Cre protein diffuses to neighboring nuclei. Cre/GFP co-immunohistochemistry revealed that only ∼56% of GFP-positive nuclei exhibited a detectable Cre signal (Fig. S1A and S1B). Given the ∼40% off-target labeling, this result suggests that Cre protein remains largely restricted to MTJ nuclei with minimal diffusion, although we cannot rule out the possibility that undetectable amounts of Cre protein contribute to leaky recombination. Next, we tested whether H2B-GFP protein is more diffusive than the NLS-sfGFP protein used earlier by performing a side-by-side comparison. Expressing H2B-GFP via MTJ-AAV resulted in a similar ∼80% labeling efficiency but a higher off-target labeling rate of ∼40% compared to ∼20% with NLS-sfGFP (Fig. S1C). These findings indicate that diffusion properties of expressed factors can vary and require careful assessment before downstream experiments.

Finally, we tested whether MTJ-AAV could be delivered systemically (Fig. 2D). Intra-orbital injection of MTJ-AAV carrying Cre-NLS into the H2B-GFP reporter successfully labeled MTJ nuclei across multiple muscles, including the gastrocnemius and diaphragm (Fig. 2E).

### Genetic labeling of MTJ ribosomes and translational profiling

Although protein synthesis is central to cellular identity and function, regionalized translation at the MTJ has not been explored, largely due to the absence of suitable tools. We hypothesized that combining MTJ-AAV with genetic ribosome tagging would enable selective profiling of ribosome-associated transcripts.

We therefore intramuscularly injected MTJ-AAV Cre-NLS or CK8-Cre-NLS into *LSL-HA-Rpl22* (RiboTag) mice (*22*), in which Cre-mediated recombination introduces an HA tag into the endogenous Rpl22 ribosomal subunit (Fig. 3A). HA immunohistochemistry confirmed the tight expression of HA-Rpl22 at the MTJ (Fig. 3B). We also performed the same experiment in Ribo-Trap mice (*Rosa26-LSL-GFP-L10a*) (*23*) and observed similar results. However, GFP-L10a displayed broader expression than HA-Rpl22 (data not shown), likely due to its overexpression from the *Rosa26* locus. For this reason, we focused on the RiboTag model for subsequent analysis. To isolate ribosome-associated transcripts, TA tissue lysates were subjected to anti-HA immunoprecipitation. Bioanalyzer profiles confirmed robust RNA recovery from Cre-delivered RiboTag muscles, with no detectable RNA in saline-injected controls (Fig. 3C). RNA sequencing was then performed to map the MTJ translational landscape (n = 4 for whole muscle, n = 3 for MTJ). Notably, MTJ markers *Tigd4* and *Col22a1* were among the most enriched ribosome-associated transcripts, validating the specificity of our dataset (Fig. 3D and 3E).

**Figure 3.**
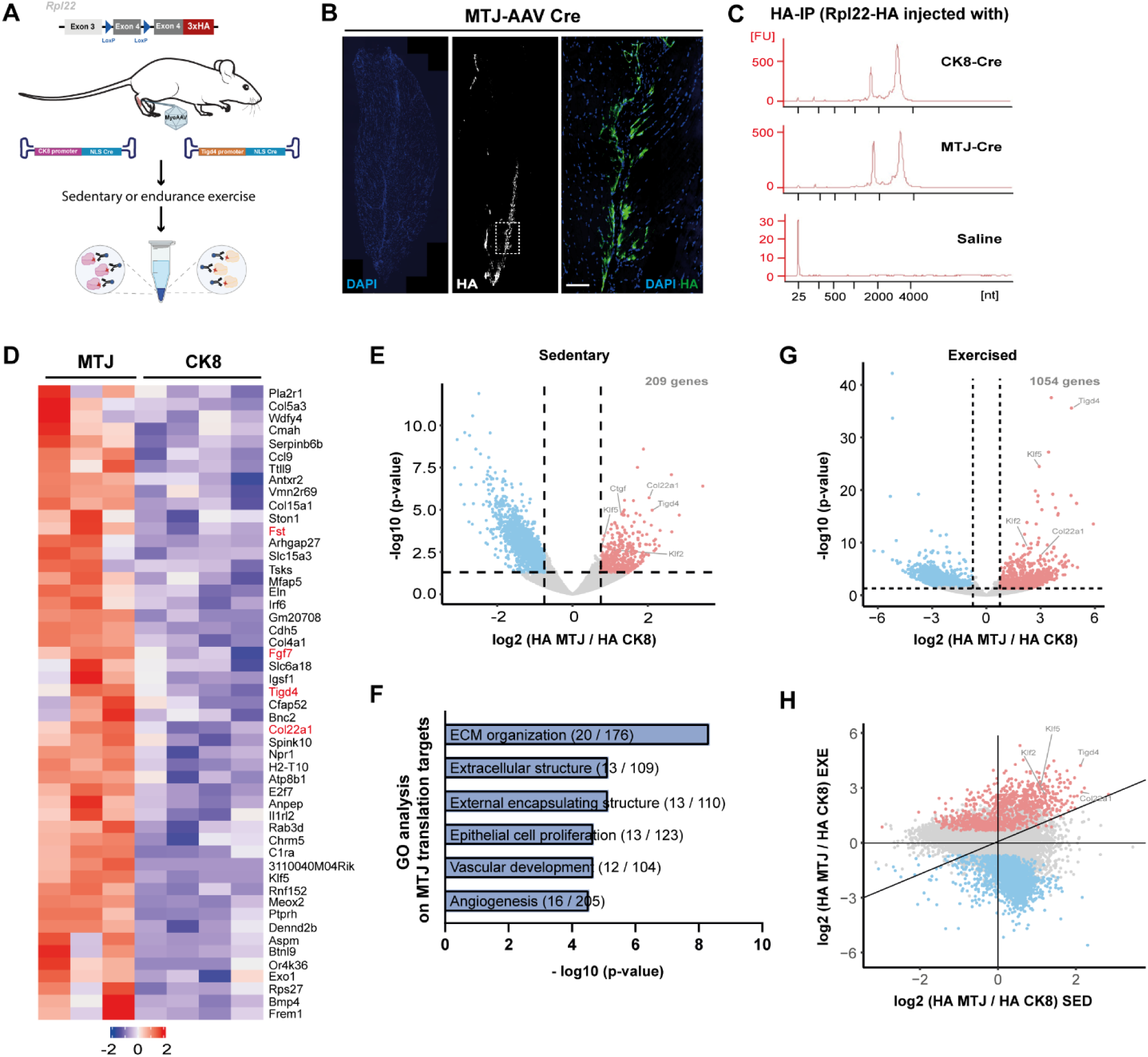
MTJ-AAV reveals a specialized translatome of MTJ. **(A)** Experimental workflow for MTJ-targeted ribosome labeling and translatome profiling in sedentary and exercised muscles. **(B)** HA immunohistochemistry confirming ribosome labeling along the MTJ. The right panel shows a magnified view of the boxed region in the middle image. Scale bar, 100 µm. **(C)** Bioanalyzer profile of RNA recovered after HA immunoprecipitation. **(D)** Heatmap showing representative MTJ-ribosome-enriched transcripts under sedentary conditions. Known transcriptional MTJ markers are indicated in red. **(E)** Volcano plot comparing CK8- and MTJ-ribosome associated transcripts in sedentary muscles. **(F)** Gene Ontology analysis of translationally enriched MTJ targets. **(G)** Volcano plot comparing CK8- and MTJ-ribosome associated transcripts following endurance exercise. **(H)** Exercise amplifies the degree of translational enrichment for MTJ-specific targets relative to sedentary conditions.

Strikingly, ∼80% of MTJ ribosome-enriched transcripts (170 of 209; fold-change > 2, p < 0.05) have not been identified as transcriptionally enriched in published snRNA-Seq datasets or in our bulk RNA-Seq from GFP-positive MTJ nuclei in Figure 1F (Fig. S2A). This indicates that they correspond to transcripts that are translationally controlled at the MTJ (Supplementary Table 1). Consistent with MTJ function, Gene Ontology analysis of these genes showed enrichment for those related to extracellular matrix biology, cytoskeletal organization, and plasma membrane structure (Fig. 3E). Intriguingly, angiogenesis-related genes were also enriched, though the significance of this remains unclear. Manual inspection identified additional targets of diverse function, including the E3 ligases *Klhl42*, *Rnf152* and *Rnf43* and transcription factors such as *Klf2* and *Klf5*. Conversely, translationally repressed genes were mainly linked to mitochondria and sarcoplasmic reticulum biology (Fig. S2B and S2C), suggesting distinct organelle composition or density at the MTJ.

### Rewiring of translational program upon sustained exercise

Endurance exercise remodels the MTJ by increasing the depth and branching of its interdigitated structures (*24*). To examine whether this event involves translational control, we subjected whole-muscle and MTJ ribosome-tagged mice to an endurance training regimen, followed by Ribosome immunoprecipitation and sequencing (Fig. 3A).

We first identified a set of genes that were translationally induced throughout the muscle in response to exercise, mainly being inflammatory mediators (*Nfkbia, Nfkb2, Stat4, Socs3*) (Fig. S2D). Notably, many of these were also induced in the MTJ translatome upon exercise, indicating a genuine muscle-wide response (Fig. S2D and S2E). Indeed, pathway analysis of genes upregulated in both CK8 and MTJ datasets revealed strong enrichment for cytokine-related signaling pathways (Fig. S2F).

We next compared MTJ ribosome-enriched transcripts to CK8-labeled ribosomes in exercised muscles (Fig. 3G). The number of MTJ-enriched genes increased substantially under exercise (1,054 versus 209 in sedentary conditions), with significant overlap between the two sets (Fig. S2G). Importantly, even shared targets exhibited markedly stronger enrichment after training (Fig. 3H). For instance, *Cpne2* increased from a 1.3-fold enrichment (log₂ scale) at rest to 2.4-fold after training, while *Klf5* rose from 1.1- to 2.9-fold (Supplementary Table 1). Together, these findings demonstrate that endurance exercise profoundly rewires the translational landscape at the MTJ, amplifying a pre-existing local translational program to meet the heightened mechanical demands of sustained activity.

### Role of untranslated regions in conferring translational specificity

Next, we asked how certain transcripts are selected for local translation. Untranslated regions (UTRs) often contain regulatory elements that control where and how efficiently mRNAs are translated (*25, 26*). To test whether such elements mediate local translation at the MTJ, we generated translation reporters based on MTJ-enriched genes. We selected *Arhgap27* and *Cpne2* as models because they were not previously identified as transcriptional markers of MTJ myonuclei but showed enrichment in our RiboTag-Seq profiling. Moreover, their UTRs are sufficiently long (∼1 kb) to harbor potential regulatory motifs yet compact enough to remain compatible with AAV packaging constraints. *Arhgap27* encodes a poorly characterized Rho GTPase–activating protein (*27*), whereas *Cpne2* encodes a similarly underexplored calcium-dependent phospholipid-binding protein that links membranes to the cytoskeleton (*28*). Both genes therefore represent plausible effectors of MTJ function, where precise coordination between cytoskeletal and membrane dynamics is essential.

We constructed AAVs in which the pan-myonuclear CK8 promoter drives an NLS-sfGFP reporter either alone or fused to the 5′ and 3′ UTRs of *Arhgap27* or *Cpne2* (Fig. 4A). In the absence of UTRs, the reporter protein was uniformly expressed throughout the myofiber, whereas inclusion of the UTRs resulted in markedly enriched expression at the fiber tips (Fig. 4B). RNAscope analysis confirmed that *GFP* transcripts remained uniformly distributed even in the presence of the UTRs (Fig. 4C), indicating that localization occurs at the translational level rather than through transcriptional control or mRNA transport. Therefore, the UTRs of *Cpne2* and *Arhgap27* are sufficient to confer MTJ-specific translation.

**Figure 4.**
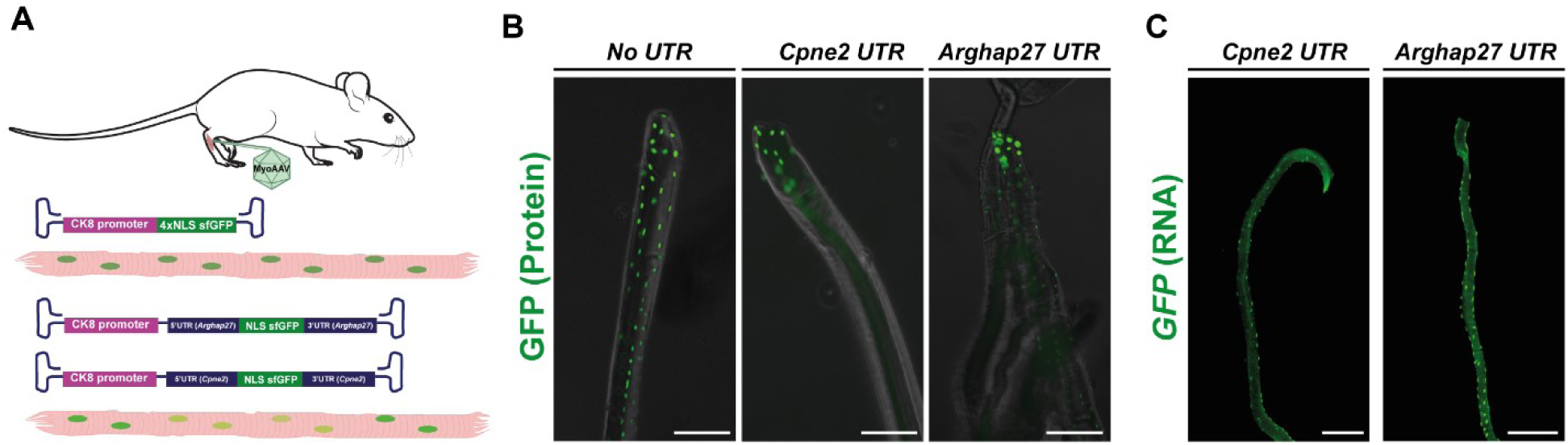
UTRs of translational target transcripts enable localized protein synthesis at the MTJ. **(A)** Schematic representation of MTJ translational reporter constructs. **(B)** Single extensor digitorum longus (EDL) myofibers were isolated one month after intramuscular injection of the AAVs shown in (A) and analyzed by epifluorescence microscopy. **(C)** EDL myofibers from (B) were fixed and subjected to RNAscope detection of GFP mRNA to assess transcript distribution. Scale bars, 100 µm.

### Translational control of KLF contributes to MTJ specific gene expression

While previous snRNA-Seq studies identified hundreds of MTJ-enriched transcripts, they did not uncover any transcription factors that are strongly and uniquely enriched at the MTJ. This contrasts with the neuromuscular junction, where key transcription factors such as Etv4 and Etv5 are specifically expressed (*6–9*). However, motif analysis of a published snATAC-Seq dataset predicted a strong enrichment of KLF-binding motifs in MTJ-specific open chromatin regions (*8*). We verified this with an independently acquired snATAC-Seq dataset (full data to be published elsewhere): 8,601 MTJ-specific chromatin accessible regions were identified that majorly located in promoters (∼50% of all peaks) (Fig. 5A). Motif analysis of the accessible regions predicted KLF factors as the top candidates (Fig. 5B). These analyses raise the question of how KLF factors might selectively function at the MTJ.

**Figure 5.**
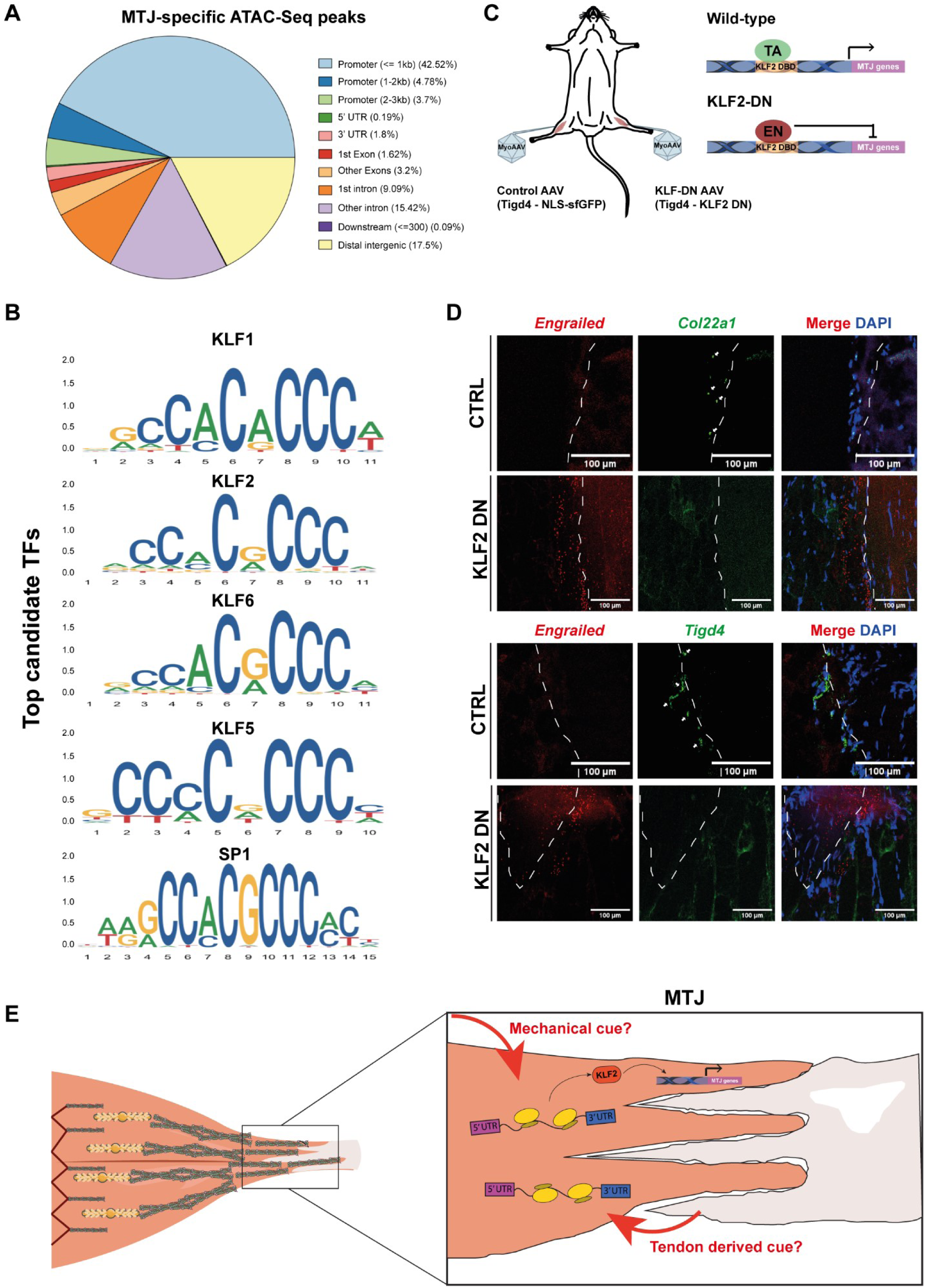
KLF transcription factors contribute to the MTJ transcriptome. **(A)** Classification of MTJ-specific ATAC-seq peaks according to their genomic locations and sequence elements. **(B)** De novo motif analysis of MTJ-specific ATAC-seq peaks identifying KLF family transcription factors as top candidates. **(C)** Schematic of KLF inhibition strategy using a dominant-negative (DN) construct expressed at the MTJ. **(D)** Expression of KLF2-DN at the MTJ suppresses MTJ marker genes *Tigd4* and *Col22a1*, as determined by RNAscope. KLF2-DN expression was detected via RNAscope targeting the *Engrailed (En)* sequence. The result was reproduced in three independent animals. **(E)** Model summarizing MTJ-specific translational regulation and its impact on local transcriptional control through KLF factors. Local signals, such as mechanical or tendon-derived cues, activate this translational program to reinforce domain specialization. Scale bars, 100 µm.

In this light, we were intrigued to find that KLF factors, *Klf2* and *Klf5*, to be enriched in our RiboTag-Seq datasets of the MTJ (Fig. 3E and 3G). We first confirmed the enrichment of Klf2 protein at the MTJ. Klf2 immunohistochemistry in tissue sections showed higher Klf2 signal in the nuclei located at the MTJ (Fig. S3A). We also surgically fractionated MTJ and non-MTJ areas using Soleus muscle. We chose this muscle type because of its flat-shaped MTJ anatomy, making the fractionation more suitable. Western blotting for Col22a1 confirmed the validity of this strategy, where Klf2 protein was also found to be more abundant at the MTJ fraction (Fig. S3B).

To probe KLF function while circumventing redundancy issue among KLF family members, we employed a dominant-negative (DN) approach. We fused the DNA-binding domain of Klf2 to the transcriptional repressor domain of *Drosophila* Engrailed (Fig. 4C), a strategy previously validated in other biological contexts (*29*). For proper comparison, we intramuscularly injected a control virus (*Tigd4* promoter expressing NLS-sfGFP) on one hindlimb, and KLF2-DN virus on the contralateral muscle (Fig. 5C). Targeted expression of Klf2-DN at the MTJ suppressed MTJ gene expression, particularly for genes containing KLF motifs identified in snATAC-Seq datasets like *Col22a1* and *Tigd4* (Fig. 5D and S3C). A reduction in Col22a1 expression was also observed at the protein level by immunohistochemistry, displaying reduced signals in KLF-DN transduced muscles (Fig. S3D). Therefore, KLF transcription factors are translationally controlled at the MTJ, where they drive a downstream transcriptional program required for MTJ specialization (Fig. 5E).

## Discussion

Our study establishes a framework for investigating regulatory mechanisms that allow the formation of specialized domains in multinucleated muscle fibers. By developing a viral system with MTJ specificity, we achieved domain-restricted genetic manipulations and ribosome tagging. These approaches uncovered extensive translational regulation at the myotendinous junction. A broad range of protein are controlled by this mechanism, encompassing extracellular matrix and cytoskeletal regulators as well as transcription factors. Previous studies largely attributed the domain formation to a transcriptional specialization of local myonuclei. The results extend our understanding of how muscle domains are formed and maintained, and expand the growing appreciation that translational control represents a central regulatory layer in cell and tissue biology. Mechanistically, our data indicate that untranslated regions of MTJ-enriched genes play a key role in conferring domain-restricted translation. Moreover, we show that the MTJ translational program is intensified by endurance exercise, indicating that this mechanism is dynamically regulated according to physiological demand. The precise cis-regulatory motifs and RNA-binding proteins involved remain to be identified. Future studies should determine how specific transcripts are selected for local translation (for example, through UTR motifs, RNA secondary structures, ribosome heterogeneity, or other mechanisms) and how these processes integrate with tissue-level signals at the MTJ such as tendon-derived or mechanical cues (Fig. 5E).

We also found that KLF transcription factors are translationally upregulated at the MTJ and contribute to local gene expression. This explains why KLF motifs are highly enriched in accessible chromatin regions of MTJ nuclei, although Klf transcript levels at the MTJ are similar to those observed in bulk myonuclei. However, KLF factors are not strictly MTJ-exclusive; they are also detectable in non-MTJ nuclei, albeit at lower levels. This suggests that KLF expression alone is insufficient to convey MTJ identity and must act in concert with additional, yet unidentified regulators to establish the full MTJ transcriptome.

A key innovation of our work is the flexibility of the MTJ-AAV strategy. Beyond transcriptional and translational profiling, it can be adapted to deliver diverse genetic tools to investigate the MTJ. Moreover, the same strategy could be extended to other muscle domains, such as the neuromuscular junction, and to other syncytial systems. For instance, distinct nuclear subtypes have been described in placental syncytiotrophoblasts by snRNA-Seq (*30*), suggesting that similar approaches could illuminate nuclear subtype-specific regulation in these contexts as well. Nonetheless, such applications must be pursued with caution, as biomolecular diffusion can confound specificity, as demonstrated by our comparisons of NLS-sfGFP versus H2B-GFP and Rpl22-HA versus GFP-L10a. With careful validation, however, MTJ-AAV and related strategies promise to become powerful tools for dissecting the spatial logic of gene regulation in syncytial cells.

## Materials and Methods

### Animals

Wild-type C57BL/6N mice were purchased from Charles River. *Rosa26-LSL-H2B-GFP* mice were described previously (*6*). RiboTag mice were obtained from Jackson Laboratory (#029977). Mice were housed under standard conditions: constant ambient temperature (23 °C), humidity (56%), and a 12-hour light/dark cycle (lights on at 6:00 am, off at 6:00 pm). Animals were euthanized by gradual CO₂ inhalation (up to 100%) over a 3-minute period. All procedures were approved by the institutional animal ethics committee of IGBMC (Comite d’Ethique). Animal health and welfare were continuously monitored by trained staff and veterinarians to minimize suffering.

### AAV construction and generation

3.3kb of *Tigd4* promoter was synthesized by GenScript and cloned into pAAV plasmid using pAAV-MCS2 (Addgene) as the backbone. Recombinant adeno-associated virus (rAAV) were generated by a triple transfection of HEK293T/17 cell line using Polyethylenimine (PEI) transfection reagent and the 3 following plasmids: the expression plasmids (pAAV), the pMyoAAV4A encoding Rep and Cap genes, and the pHelper (Agilent) encoding the adenovirus helper functions. 48 h after transfection, rAAV vectors were harvested from cell lysate and treated with Benzonase (Merck) at 100U/mL. They were further purified by gradient ultra-centrifugation with Iodixanol (OptiprepTM density gradient medium) followed by dialysis and concentration against Dulbecco’s Phosphate Buffered Saline (DPBS) using centrifugal filters (Amicon Ultra-15 Centrifugal Filter Devices 100K, Millipore). Viral titres were quantified by Real-Time PCR using the LightCycler480 SYBR Green I Master (Roche) and primers targeting respective insert sequences (*e.g.,* sfGFP). Titers are expressed as genome copies per milliliter (GC/mL). The pMyoAAV4A plasmid was self-constructed by IGBMC’s molecular biology platform (*31*), benchmarking the published construct (*21*).

### AAV injection

Mice were anesthetized via intraperitoneal injection of a ketamine/xylazine mixture composed of 10% ketamine (100 mg/mL), 5% xylazine (20 mg/mL), and 85% sterile NaCl, administered at a total dose of 100 µL per 10 g of body weight. Anesthesia depth was verified by the absence of pedal reflex before proceeding with the injections.

For intramuscular (IM) injection, the skin over the target muscle was disinfected with 70% ethanol under anesthesia. MyoAAV4a vectors carrying different constructs were injected intramuscularly into the tibialis anterior muscle using a 30-gauge needle. Each construct was administered at its respective final viral genome (vg) dose per muscle: 5 × 10¹⁰ - 10¹¹ vg, depending on the viral concentration and experimental design. The total injection volume per muscle was adjusted with sterile NaCl to ensure a constant final volume of 20 µL across all conditions. Contralateral muscles received vehicle or a control vector where applicable. Muscles were collected at 1-month post-injection for molecular and histological analyses.

For systemic administration, anesthetized mice were placed in a prone position, and MyoAAV4a particles were delivered via retro-orbital sinus injection using a 30-gauge insulin syringe. A total volume of 100 µL corresponding to 1.5 x 10^13^ vg per mouse was injected slowly over approximately 15 seconds to minimize reflux. Animals were monitored until full recovery and returned to their cages. Tissue collection was performed at 2 months post-injection.

### Ribosome immunoprecipitation and sequencing (RiboTag-Seq)

10 weeks old homozygous RiboTag mice were intramuscularly injected with CK8- or MTJ-AAV carrying Cre recombinase. TA muscles were harvested one month later, snap-frozen, and stored at –80°C until processing.

Frozen tissues were thawed on ice and transferred to homogenization tubes containing silica beads (Precellys P000918-LYSK1-A). For the MTJ-AAV condition, lysates were pooled from four injected muscles from different mice. For the CK8-AAV condition, lysates were pooled from two injected muscles and two un-injected WT muscles to normalize total lysate concentration between conditions, as this might affect HA-based immunoprecipitation efficiency and purity.

Each sample was homogenized in 1 ml lysis buffer containing 50 mM Tris-HCl (pH 7.5), 100 mM KCl, 12 mM MgCl₂, 1% NP-40, 1 mM DTT, 200 U/ml RNAsin (Promega), 1 mg/ml heparin (Sigma), 100 µg/ml cycloheximide (Sigma), and protease inhibitor cocktail (Roche). DTT, RNAsin, heparin, cycloheximide and protease inhibitor were added freshly before use. Tissues were minced with sterile scissors and incubated on ice for 20 minutes before homogenization using the Precellys Evolution system (5,000 rpm, 25 seconds, twice, with 10-minute intervals on ice). Lysates were clarified by centrifugation at 10,000g for 20 minutes at 4°C, and the supernatants were transferred to fresh tubes.

Protein concentration was determined via Bradford assay (Bio-Rad). Approximately 3 mg of lysate was incubated with 5 µl HA antibody (Covance, clone 12CA5) for 4 hours at 4°C with gentle rotation. Meanwhile, magnetic Protein A/G beads (Pierce) were equilibrated in lysis buffer. Forty microliters of beads were added to each sample, followed by overnight incubation at 4°C with rotation.

Beads were washed three times with high-salt buffer (same as lysis buffer, but with 300 mM KCl and 0.5 mM DTT), each wash involving 5 minutes of rotation. After the final wash, beads were resuspended in 800 µl high-salt buffer and transferred to a new tube. RNA was extracted by adding 350 µl of RLT buffer (Qiagen RNeasy kit) and following the manufacturer’s protocol, including DNase treatment. Purified RNA was stored at –80°C until further use.

Library preparation was performed at the GenomEast platform at the Institute of Genetics and Molecular and Cellular Biology using Illumina Stranded Total RNA Prep Ligation with Ribo-Zero Plus (Reference Guide - PN 1000000124514). Total RNA-Seq libraries were generated from 50 ng of total RNA using Illumina Stranded Total RNA Prep, Ligation with Ribo-Zero Plus kit and IDT for Illumina RNA UD Indexes, Ligation (Illumina, San Diego, USA), according to manufacturer’s instructions. DNA library were amplified using 14 cycles of PCR. Surplus PCR primers were further removed by two successive purifications using SPRIselect beads (Beckman-Coulter, Villepinte, France). The final libraries were checked for quality and quantified using Bioanalyzer 2100 system (Agilent technologies, Les Ulis, France). Libraries were sequenced on an Illumina NextSeq 2000 sequencer as paired-end 50 base reads. Image analysis and base calling were performed using RTA version 2.7.7 and BCL Convert version 3.8.4.

### Endurance exercise

RiboTag mice were intramuscularly injected with CK8-Cre-NLS or Tigd4-Cre-NLS AAV. One month post-injection, the mice underwent an endurance exercise regimen using an animal treadmill (LE8710, Panlab, Harvard Apparatus, Spain). To ensure proper acclimatization, the mice were introduced to the treadmill environment over a period of three days prior to the start of the experimental protocol. During this familiarization phase, the treadmill operated at a low speed of 10–12 cm/s for 15 minutes each day. The exercise training protocol began at 12 cm/s for 15 minutes, increased to 20 cm/s for 50 minutes, and finished with a 5-minute cooldown at 10 cm/s. All exercise sessions were consistently conducted at the same time of day for 4 weeks (5 days consecutively and 2 days rest).

### FACS isolation and low-input RNA-Seq

TA muscles injected with CK8-NLS-sfGFP or Tigd4-NLS-sfGFP AAVs were collected, snap frozen, and stored at –80°C until further use. The TA muscles were minced and incubated in 300µl of cold hypotonic buffer (250mM sucrose, 10mM KCl, mM MgCl2, 10mM Tris-HCl pH 8, 25mM HEPES pH 8, 0.3% Triton x-100, 0.2mM PMSF, 0.1 mM DTT, and 0.2U/µL RNase inhibitor) for 5 mins. Samples were then transferred to 2ml ‘Tissue homogenizing CKMix’ (Bertin Technologies) with an additional 700µl hypotonic buffer. Following a further 15mins incubation, the samples were then homogenized with the ‘Precellys 24 tissue homogenizer’ (Bertin Technologies) for 25s at 5,000rpm. The homogenized samples were then passed through a 100µm filter (Sysmex) followed by a 20µm filter (Sysmex) before the nuclei were pelleted by centrifugation at 400g for 10 min at 4 °C. The pellets were then washed and resuspended in a washing buffer (2% BSA in PBS + RNase inhibitor 0.2U/µL). The centrifugation and wash steps were then repeated before the homogenized samples were passed through a ‘5ml polystyrene round-bottom tube with cell-strainer cap’ (Corning). DAPI was then added to the resuspended nuclei at a final concentration of 200mM. The isolated nuclei were sorted with FACs Aria Fusion 2022, with BD FACSDiva and FlowJo (v10) software, to sort out the DAPI and GFP+ nuclei. Approximately 1,000 nuclei were collected from each sample. The sorted nuclei were immediately collected in 11.5µL of CDS sorting solution, SMART-Seq® mRNA LP (with UMIs) recipe (Takara), briefly spun down, then flash-frozen on dry ice. The frozen isolated nuclei were then stored at -80°C until further use.

Library preparation was performed at the GenomEast platform at the Institute of Genetics and Molecular and Cellular Biology using Takara Bio USA, Inc., SMART-Seq mRNA User Manual - PN 062223 + Illumina Nextera XT DNA Library Prep Kit (Reference Guide - PN 15031942). Full length cDNA were generated from 100 to 2000 nuclei using SMART-Seq mRNA (Takara Bio Europe, Saint Germain en Laye, France) according to manufacturer’s instructions with 11 cycles of PCR for cDNA amplification by Seq-Amp DNA polymerase. The entire volume of each pre-amplified cDNA was then used as input for Tn5 transposon tagmentation followed by 12 cycles of library amplification using Nextera XT DNA Library Preparation Kit and IDT for Illumina DNA/RNA UD Indexes, Tagmentation (Illumina, San Diego, USA). Following purification with SPRIselect beads (Beckman-Coulter, Villepinte, France), the size and concentration of libraries were assessed by capillary electrophoreris (Bioanalyzer 2100 system, Agilent technologies, Les Ulis, France). Full length cDNA was generated from from 100 to 2000 nuclei using SMART-Seq mRNA Kit (Takara Bio Europe, Saint Germain en Laye, France) according to manufacturer’s instructions with 11 cycles of PCR amplification by Seq-Amp polymerase. Totality of pre-amplified cDNA were then used as input for Tn5 transposon tagmentation followed by 12 cycles of library amplification using Nextera XT DNA Library Preparation Kit and IDT for Illumina DNA/RNA UD Indexes (Illumina, San Diego, USA). Following purification with SPRIselect beads (Beckman-Coulter, Villepinte, France), the size and concentration of libraries were assessed using Bioanalyzer 2100 system (Agilent technologies, Les Ulis, France). Libraries were sequenced on an Illumina NextSeq 2000 sequencer as paired-end 50 base reads. Image analysis and base calling were performed using RTA version 2.7.7 and BCL Convert version 3.8.4.

### Bioinformatic analyses of RNA-Seq datasets

All sequencing was performed on an Illumina NextSeq 2000 platform in a 2x50bp paired-end configuration. For RiboTag-Seq, sequencing data was processed processed using PiGx-RNA-seq (30277498) pipeline. In short, the data was mapped onto the GRCm39/mm11 version of the mouse transcriptome (downloaded from the ENSEMBL database 29155950) using SALMON (28263959). The quantified data was processed using tximport (26925227), and the differential expression analysis was done using DESeq2 (25516281). Genes with less than 5 reads in all biological replicates of one condition were filtered out before the analysis. Two groups of differentially expressed genes were defined - a relaxed set containing genes with an absolute log2 fold change of 0.5, and a stringent set containing games with an absolute log2 fold change of 1. The fold change was deemed significant if the adjusted p-value was less than 0.05 (Benjamini-Hochberg corrected). For bulk RNA-Seq, reads were preprocessed with cutadapt 4.2 to remove adaptor sequences, poly(A) tails, and low-quality bases. Reads shorter than 40 bp were discarded. The remaining reads were mapped to Mus musculus rRNA sequences using bowtie2 2.3.5 (*32*) and reads aligning to these sequences were removed. The filtered reads were then aligned to the Mus musculus reference genome (GRCm39 assembly) using STAR 2.7.10b (*33*). Gene-level quantification was performed with HTSeq-count 1.99.2 (*34*) in “union” mode, using Ensembl 111 annotations. Differential gene expression analysis was carried out with the DESeq2 1.34.0 (*35*) R/Bioconductor package using default parameters. P-values were adjusted for multiple testing with the Benjamini-Hochberg method.

Gene ontology and pathway reactome analyses were performed using ‘https://maayanlab.cloud/Enrichr/’ website.

### Analyses of snATAC-Seq dataset and motif enrichment

Single-nucleus ATAC-seq (snATAC-seq) libraries were processed using Signac v1.10.0 and Seurat v4.3.0 in R. Raw peak files were imported and converted to GRanges objects for genomic coordinate standardization. Cell-type-specific open chromatin peaks were identified for myotendinous junction (MTJ) nucleus through Seurat object integration with parallel transcriptomic data (GEX), utilizing clustering and differential accessibility analysis to isolate peaks enriched in MTJ-specific clusters. From the total accessible chromatin landscape, myotendinous junction-specific peaks were extracted by filtering for regions showing significant accessibility exclusively in MTJ nucleus relative to other muscle nucleus populations. Peak coordinates were standardized using StringToGRanges coordinate conversion (mm10 genome build). Selected peaks were then used as input for downstream transcription factor binding site discovery via motif scanning with JASPAR 2020 vertebrate PWMs.

Transcription factor binding site (TFBS) enrichment analysis was performed on MTJ-specific open chromatin regions using a computational motif-scanning approach. MTJ-specific accessible chromatin peaks were identified from ATAC-seq data and used as input for systematic motif discovery. Vertebrate transcription factor position weight matrices (PWMs) were obtained from the JASPAR 2020 database using TFBSTools (v1.38.0). All vertebrate PWMs with non-redundant versions were retrieved, encompassing 746 motif models representing diverse transcription factor families. Genomic sequences corresponding to each peak region were extracted from the mouse reference genome (mm10) using BSgenome.Mmusculus.UCSC.mm10 (v1.4.3). Motif scanning was performed with motifmatchr (v1.22.0) using the matchMotifs() function, which implements a log-odds scoring algorithm based on PWM models. For each peak-motif pair, a match score was computed by sliding the PWM across both DNA strands. Binding sites exceeding the default log-odds threshold (typically p < 10⁻⁴) were classified as putative TFBS occurrences.

Results were integrated into a Seurat object (v4.3.0) using the Signac framework (v1.10.0) for chromatin accessibility data. A binary motif occurrence matrix was constructed with dimensions n peaks × m motifs, where each entry indicates presence (1) or absence (0) of a motif within a given peak. This matrix was stored as a ChromatinAssay object within the Seurat framework, enabling downstream integration with single-cell chromatin accessibility profiles when applicable.

Global motif enrichment was calculated as the proportion of peaks containing each motif: Enrichment (%) = (Number of peaks with motif / Total peaks) × 100 Motifs were ranked by frequency of occurrence across all analyzed peaks compared to random occurences. The top-ranking motifs representing the most enriched transcription factor binding sites in MTJ-specific open chromatin were identified for downstream validation and functional interpretation.

MTJ-specific accessible chromatin peaks were functionally annotated using ChIPseeker v1.36.0 with the mouse genome annotation database (TxDb.Mmusculus.UCSC.mm10.knownGene and org.Mm.eg.db). Peak coordinates were mapped to their nearest genomic features, including promoters (±3 kb from transcription start sites), gene bodies, introns, and intergenic regions, using annotatePeak(). The distribution of peaks across genomic features was visualized with annotation pie charts to assess regulatory landscape composition. Gene symbols associated with each peak were extracted, and genes with multiple associated peaks (≥150 peaks per gene) were identified as high-confidence MTJ regulatory targets. This threshold-based filtering enriched for genes with extensive regulatory architecture characteristic of master regulators and structural genes defining myotendinous junction identity.

All analyses were performed in R (v4.3.1). The complete analysis pipeline, including motif occurrence matrices and enrichment statistics, was saved for reproducibility and downstream chromVAR-based activity inference.

### Histology and imaging

For histological analysis, samples were rapidly embedded in Cryomatrix (Epredia) in cryomolds then snap-frozen in isopentane pre-cooled in liquid nitrogen for approximately 15 seconds until the cryomatrix solidified completely. Samples embedded in cryomatrix were equilibrated to cryostat temperature (–20 °C) prior to sectioning. Transverse cryosections were cut at 10 μm thickness using a Leica CM3050S cryostat (Leica Microsystems, Germany) and collected on Superfrost Plus adhesion microscope slides (Epredia). Sections were then stored at –20 °C for 30 mins then stored at -80 °C until further use.

For RNAscope, the tissue sections were fixed with 4% PFA in PBS for 15 minutes at 4 °C. After two times washing with PBS, the sections were serially dehydrated through increasing ethanol (50%, 70%, 100%) 5 minutes each. Subsequently, RNAscope was performed according to manufacturer’s guideline using the RNAscope Multiplex Fluorescent Reagent Kit v2 (Bio-Techne 323100). Proteinase IV was used for our procedure. The following RNAscope probes were used for our study: *Col22a1* (590911-C2), *Tigd4* (598761), *Ttn* (483031), *GFP* (409011) and *Engrailed* (newly designed for this study). After all RNAscope procedures, samples were counterstained with DAPI and mounted using Prolong Gold Antifade (Thermofisher scientific). When combined with antibody staining, tissue sections were processed to blocking buffer (see below) after the last wash and proceeded to regular immunohistochemistry.

For immunofluorescence staining, transverse cryosections were fixed in 4% paraformaldehyde (PFA) for 10 minutes at room temperature (RT), followed by three washes in PBT (PBS containing 0.1% Tween-20) for 5 minutes each. Permeabilization was performed in PBX (PBS containing 0.5% Triton X-100) for 6 minutes, after which sections were rinsed in PBS for 5 minutes. Non-specific binding was blocked by incubating the sections in blocking buffer (PBS containing 5% BSA, 3% horse serum and 0.1% Triton X-100) for 1 hour at RT. Sections were then incubated with primary antibody diluted in blocking buffer overnight at 4 °C in a humidified chamber. The following day, slides were washed three times in PBT for 10 minutes each, then incubated with fluorophore-conjugated secondary antibody diluted in blocking buffer for 1 hour at RT. DAPI (1 μg/mL) was included in the secondary incubation step for nuclear counterstaining. Finally, sections were washed three times in PBT for 10 minutes each in the dark, mounted with ProLong Gold Antifade, and stored at 4 °C until imaging. The following antibodies were used in this study: GFP (Aves Labs; 1:500), Klf2 (Cell Signaling, 1:500), Dystrophin (Abcam, 1:250), and Col22a1 (Gift from Manuel Koch, 1:1000).

Images were acquired on a Leica confocal at the IGBMC imaging core facility. 20× oil-immersion objective was used. Fluorescent signals were sequentially detected using appropriate laser lines and emission filters for DAPI, Opal 520, Opal 570 and Opal 690 (Opal chemicals were from Akoya Biosciences) to avoid channel bleed-through. Laser power, gain, and offset were optimized and maintained constant across samples. Imaging was performed using the tile scan mode to capture large tissue areas, with a frame size of 520 × 520 pixels and processed using imageJ.

### Western blotting

TA tissues were lysed as described in the ‘Ribosome immunoprecipitation and sequencing’ section. Lysates were denatured by adding Laemmli buffer and boiling for 10 minutes. Denatured samples were separated by SDS-PAGE (Bio-Rad) and transferred into nitrocellulose membrane (Amarsham). Transferred membranes were blocked for one hour with 5% skim milk in TBS plus 0.1% Tween-20 (TBST) at RT. Afterwards, primary antibodies were incubated in 5% BSA in TBST supplemented with 0.1% sodium azide overnight in cold room with gentle rocking. After three times of washing with TBST (10 minutes each), membranes were incubated with secondary antibodies (anti-mouse or anti-rabbit HRP; Cell Signaling) diluted in skim milk (1:5000) for one hour in RT. After three times washing with TBST, membranes were developed using chemiluminescence (ECL substrate; Pierce) and Amarsham ImageQuant 800. For Streptavidin-HRP, antibody was incubated for one hour in skim milk after blocking.

The following antibodies were used in this study: Col22a1 (Abcam; 1:500), β-actin (Cell Signaling; 1:1000), Klf2 (Cell Signaling, 1:500), HA (Covance, 1:2000), and Streptavidin-HRP (Sigma, 1:5000).

### Statistical analysis

Graph generation and statistical analyses were performed using GraphPad Prism as described in each figure legend.

## Supporting information

RiboTag-Seq marker genes

## Acknowledgments

## Funding

We thank grant supports to M.K. from the

European Research Council ERC-StG 101039531 (M.K)

ANR-22-CE13-0023-03 MYODOM (M.K)

ANR t-ERC StG 2021 (M.K) LABEX INTR (M.K)

University of Strasbourg IDEX Attractivitae (M.K)

INSERM ATIP-Avenir (M.K)

AFM-Telethon Trampoline 24287 and n°22AA003-00 (M.K)

Grand Est PhD fellowship (N.E.K)

Fondation pour la Recherche Médicale postdoctoral fellowship (S.M)

AFM n°28842 (P.M)

ANR-21-CE14-0042-01 MOTOMYO (P.M)

## Author contributions

Conceptualization: M.K

Main Experiments: J.N and N.E.K

Other Experiments: C.S, S.M, and L.Y

Bioinformatic analyses: V.F and E.J

Supervision: P.M and A.A

Methodology (AAV generation): E.L

Writing: M.K

## Data and materials availability

All data and materials used for this study are available from the corresponding author upon reasonable request. All NGS datasets have been deposited to EBI (RNA-Seq: E-MTAB-16305; RiboTag-Seq under sedentary: E-MTAB-16304; RiboTag-Seq after exercise: E-MTAB-16306). The codes used in this study are available in Github server (https://github.com/BIMSBbioinfo/MTJ_Figures). All data are available in the main text or the supplementary materials.

