## Supplementary data for "Local translational program at the muscle-tendon junction endows domain identity in muscle syncytia"

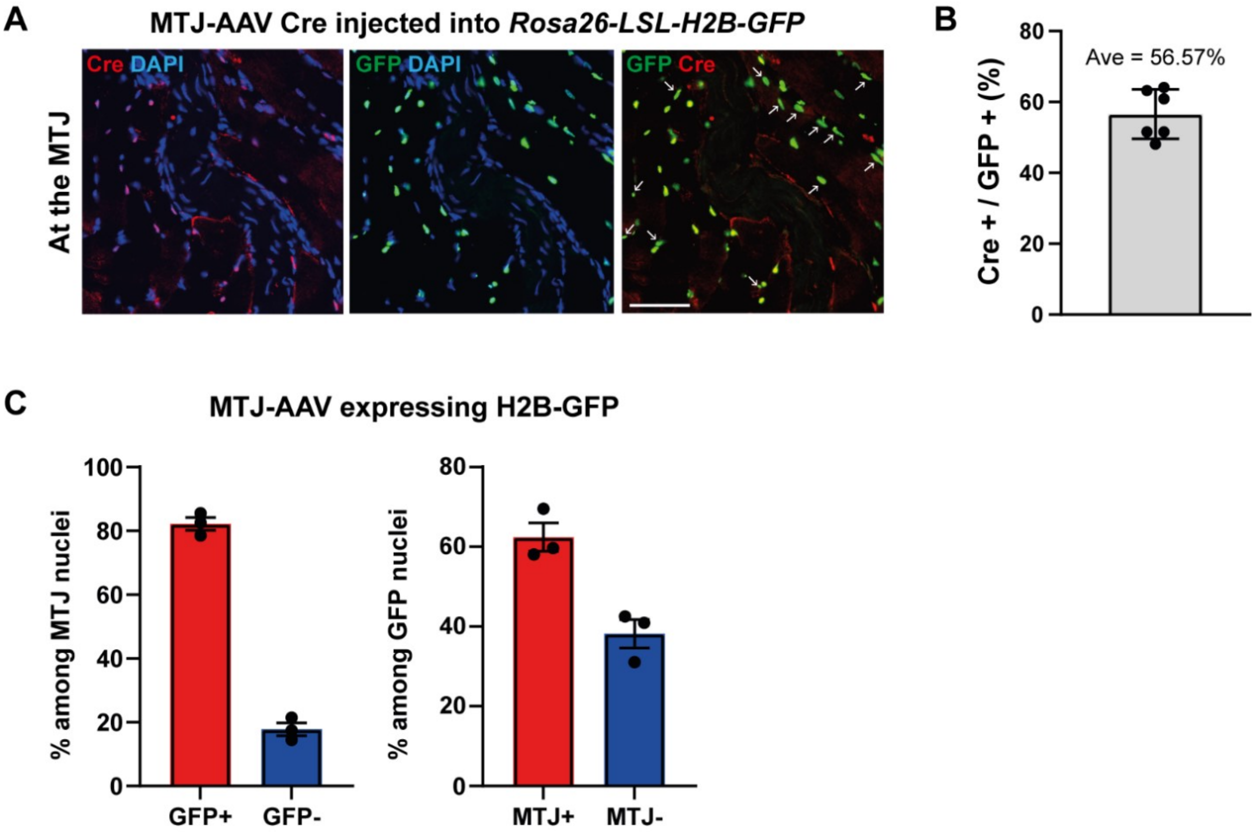


**Figure S1. H2B-GFP is more diffusive than 4xNLS-sfGFP**

**(A)** Immunohistochemistry for Cre and GFP in TA muscles of *Rosa26-LSL-H2B-GFP* mice injected with MTJ-AAV-Cre-NLS. Arrows indicate GFP-positive but Cre-negative nuclei. Scale bars, 50 μm.

**(B)** Quantification of Cre-positive nuclei among GFP-expressing nuclei (n = 6).

**(C)** Wild-type mice were intramuscularly injected with MTJ-AAV-H2B-GFP, and labeling efficiency and specificity were quantified (n = 3).

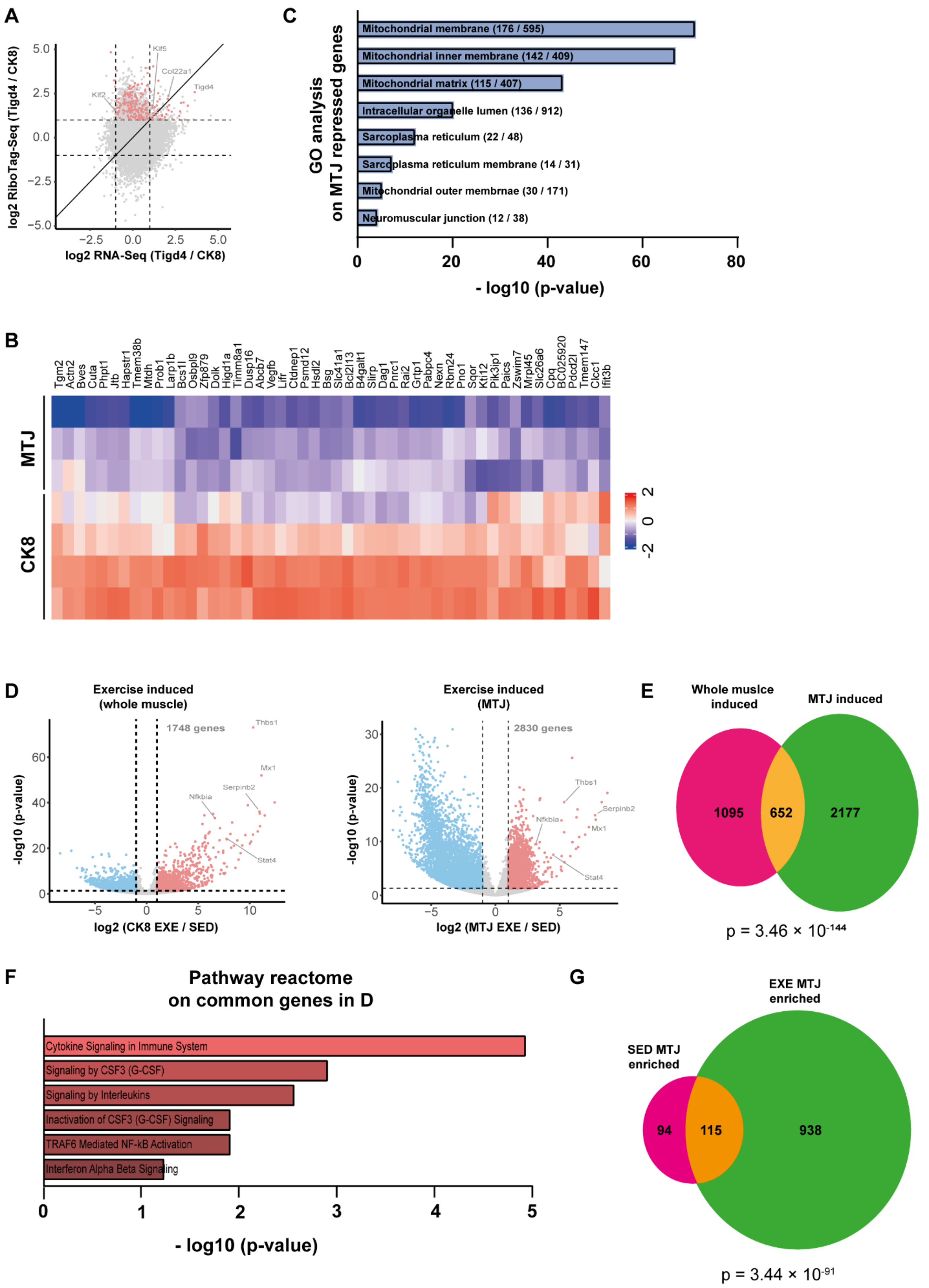

A
log2 RiboTag-Seq (Tigd4 / CK8)
log2 RNA-Seq (Tigd4 / CK8)
Klf5
Col22a1
Tigd4
Klf2
C
GO analysis on MTJ repressed genes
Mitochondrial membrane (176 / 595)
Mitochondrial inner membrane (142 / 409)
Mitochondrial matrix (115 / 407)
Intracellular organelle lumen (136 / 912)
Sarcoplasma reticulum (22 / 48)
Sarcoplasma reticulum membrane (14 / 31)
Mitochondrial outer membrnae (30 / 171)
Neuromuscular junction (12 / 38)
- log10 (p-value)
B
Tgm2 Actn2 Bves Cuta Phpt1 Jtb Hapstr1 Tmem38b Mtdh Prob1 Larp1b Bcs1l Osbpl9 Zfp879 Dolk Higd1a Timm8a1 Dusp16 Abcb7 Vegfb Lifr Ctdnep1 Psmd12 Hsdl2 Bsg Slc41a1 Bcl2l13 B4galt1 Slirp Dag1 Pnrc1 Rai2 Grtp1 Pabpc4 Nexn Rbm24 Pno1 Sqor Ktl12 Pik3ip1 Paics Zswim7 Mrpl45 Slc26a6 Cpq BC025920 Pdcd2l Tmem147 Clcc1 Ifit3b
MTJ
CK8
D
Exercise induced (whole muscle)
1748 genes
Thbs1
Mx1
Serpinb2
Nfkbia
Stat4
-log10 (p-value)
log2 (CK8 EXE / SED)
Exercise induced (MTJ)
2830 genes
Thbs1
Nfkbia
Serpinb2
Mx1
Stat4
-log10 (p-value)
log2 (MTJ EXE / SED)
E
Whole muslce induced
MTJ induced
1095
652
2177
p = 3.46 × 10^-144
F
Pathway reactome on common genes in D
Cytokine Signaling in Immune System
Signaling by CSF3 (G-CSF)
Signaling by Interleukins
Inactivation of CSF3 (G-CSF) Signaling
TRAF6 Mediated NF-kB Activation
Interferon Alpha Beta Signaling
- log10 (p-value)
G
EXE MTJ enriched
SED MTJ enriched
94
115
938
p = 3.44 × 10^-91

**Figure S2. Analyses related to MTJ translational profiling**

**(A)** Comparison of fold-enrichments between RNA-Seq in Figure 1F and RiboTag-Seq. The colored genes are those identified as up-regulated in Figure 3E.

**(B)** Heatmap showing representative MTJ ribosome-depleted transcripts.

**(C)** Gene Ontology analysis of translationally depleted genes in the MTJ.

**(D)** Comparison of genes enriched in whole-muscle ribosomes (CK8-Cre–labeled) or MTJ ribosomes (Tigd4-Cre–labeled) following exercise relative to sedentary conditions.

**(E)** Venn diagram showing overlap between translationally upregulated genes upon exercise in whole muscle and MTJ. The p-value was calculated using a hypergeometric test with a total background of 21,900 protein-coding genes in the mouse genome.

**(F)** Reactome pathway analysis of the commonly enriched genes identified in (D).

**(G)** Venn diagram showing overlap between translationally upregulated genes at the MTJ under sedentary and exercised conditions. The p-value was calculated using a hypergeometric test with a total background of 21,900 protein-coding genes in the mouse genome.

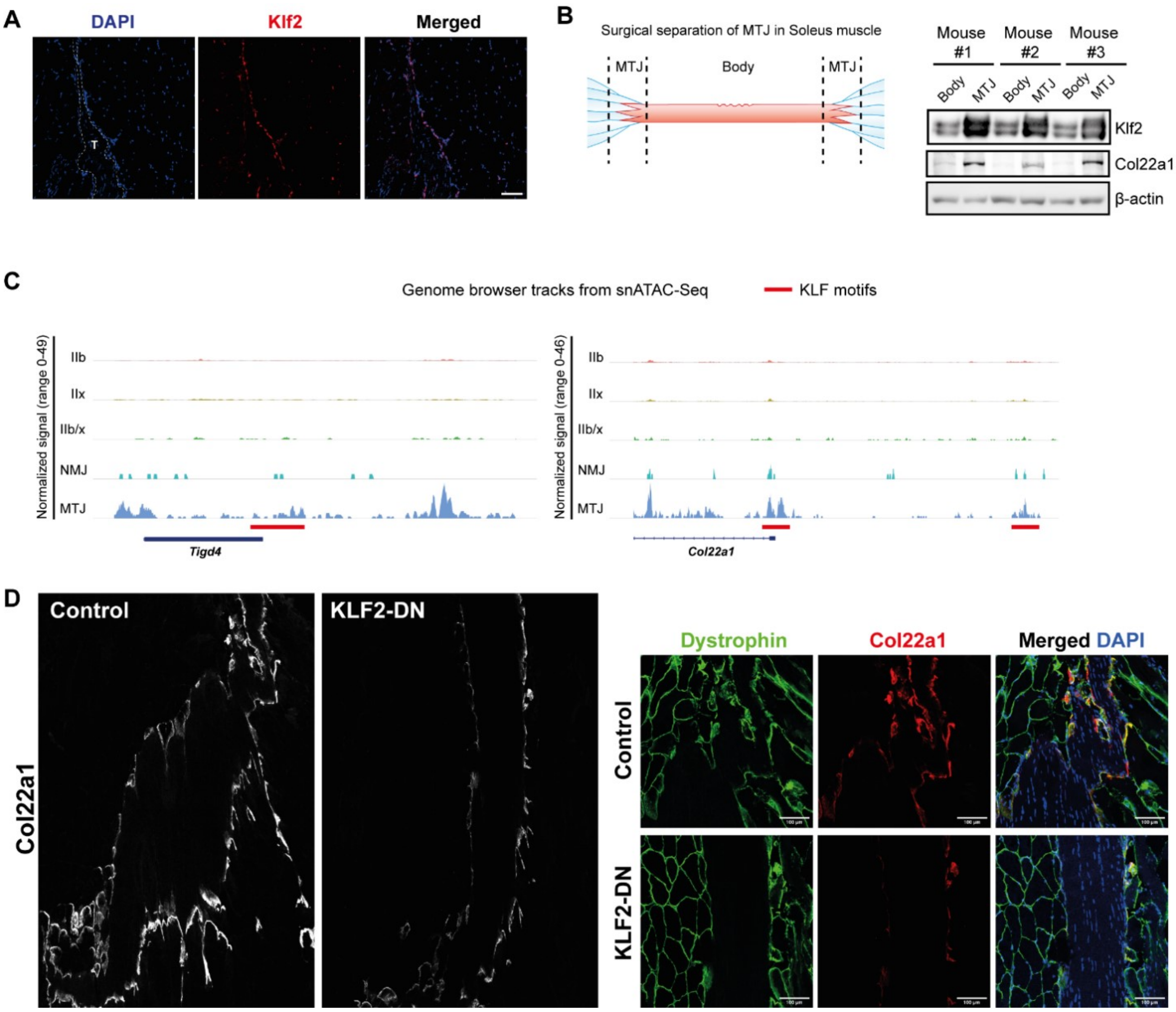


**Figure S3. Expression of KLF2 protein at the MTJ**

**(A)** Longitudinal sections of TA muscle were immunostained with an anti-KLF2 antibody. Note the stronger expression of KLF2 along the MTJ. T, tendon.

**(B)** MTJ and non-MTJ parts of Soleus muscle were surgically separated and examined by Western blotting (n=3).

**(C)** Genome browser tracks of ATAC-Seq peaks at the *Tigd4* and *Col22a1* genes and location of KLF motifs.

**(D)** Reduction of Col22a1 protein expression at the MTJ of KLF2-DN injected muscles. Left images are tile scan of entire MTJ area, and the right images are magnification of subparts. Note that Dystrophin expression is similar, while only Col22a1 expression is decreased by KLF2-DN. The result was reproduced in three independent animals.

Scale bars, 100 μm.

**Table S1. Marker genes from RiboTag-Seq experiments**

Sheet #1: Translationally induced genes in whole muscle (CK8) after exercise

Sheet #2: Translationally induced genes in MTJ after exercise

Sheet #3: Translationally enriched genes at the MTJ compared to whole muscle in sedentary condition

Sheet #4: Translationally enriched genes at the MTJ compared to whole muscle after exercise
